# Reduced protein kinase C delta association with a higher molecular weight complex in mitochondria of Barth Syndrome lymphoblasts

**DOI:** 10.1101/2021.07.21.453087

**Authors:** Edgard M. Mejia, Hana M. Zegallai, Genevieve C. Sparagna, Grant M. Hatch

## Abstract

The protein kinase C delta (PKCδ) signalosome exists as a high molecular weight complex in mitochondria and controls mitochondrial oxidative phosphorylation. Barth Syndrome (BTHS) is a rare X-linked genetic disease in which mitochondrial oxidative phosphorylation is impaired due to a mutation in the gene TAFAZZIN which results in reduction in the phospholipid cardiolipin and an accumulation of monolysocardiolipin. Here we examined if PKCδ association with a higher molecular weight complex was altered in mitochondria of BTHS lymphoblasts. Immunoblot analysis of blue native-polyacrylamide gel electrophoresis mitochondrial fractions revealed that PKCδ associated with a higher molecular weight complex in control lymphoblasts but this was markedly reduced in BTHS patient B lymphoblasts in spite of an increase in PKCδ protein expression. We hypothesize that the lack of PKCδ within this higher molecular weight complex may contribute to defective mitochondrial PKCδ signaling and thus to the bioenergetic defects observed in BTHS.

## Introduction

Barth Syndrome (BTHS) is a rare X-linked genetic disease caused by a mutation in the TAFAZZIN gene localized on chromosome Xq28.12 [1-3]. BTHS is characterized by cardiomyopathy, skeletal myopathy, growth retardation, neutropenia, and frequently 3-methylglutaconic aciduria. The TAFFAZIN gene encodes the protein tafazzin. Tafazzin is a transacylase enzyme involved in the remodeling of the mitochondrial phospholipid cardiolipin (CL) [4,5]. BTHS mutations in TAFAZZIN result in reduced CL and elevated monolysocardiolipin [1-3, 6]. In several studies Epstein-Barr virus transformed patient lymphoblasts have been used to examine BTHS metabolic pathology [6-9].

Protein kinase C delta (PKCδ) is a signaling kinase that regulates many cellular responses and is controlled via multi-site phosphorylation [10-13]. The PKCδ pathway adjusts the fuel flux from glycolytic sources to the intensity of mitochondrial respiration, thus controlling mitochondrial oxidative phosphorylation. In mitochondria, the PKCδ signalosome exists in a high molecular weight complex, which includes cytochrome *c* as the upstream driver of PKCδ, the adapter protein p66Shc as a platform and retinol [12, 14]. All four components are required for activation of PKCδ signaling in mitochondria. The expression of PKCδ and its association with a higher molecular weight complex in BTHS cells had never been investigated. We previously demonstrated that PKCδ phosphorylation was altered on several sites in BTHS patient lymphoblasts [15]. Since altered phosphorylation may affect PKCδ activation, we examined if PKCδ association with a higher molecular weight complex was altered in mitochondria of BTHS lymphoblasts.

In this study we demonstrate for the first time that PKCδ is associated with a higher molecular weight complex in lymphoblast mitochondria and that this is reduced in BTHS patient B lymphoblast mitochondria compared to age-matched control lymphoblasts in spite of an increase in PKCδ protein expression. We hypothesize that the lack of PKCδ within this complex may contribute to defective mitochondrial PKCδ signaling and thus to the bioenergetic defects observed in BTHS.

## Materials and Methods

Epstein-Barr virus transformed BTHS lymphoblasts (Patient 618: Exon 2, c. 171 del. A, frameshift) and age-matched control lymphoblasts (Patient 3798) were a generous gift from Dr. Richard Kelley, Kennedy Kreiger Institute, Baltimore, MD., and cultured as previously described [9]. Electrospray ionisation mass spectrometry (ESI-MS) coupled with high performance liquid chromatography (HPLC) mass spectrometry of cardiolipin (CL) and monolysocardiolipin (MLCL) from cell lysates was performed described [16]. Mitochondrial fractions were isolated using the mitochondrial isolation kit from Abcam (Toronto, ON, Canada, Catalog number ab110170). Mitochondrial protein content was determined using the M protein assay kit (Mississauga, ON, Canada). For Blue-Native polyacrylamide gel electrophoresis (BN-PAGE) analysis, mitochondrial protein (80 µg) was treated with 0.2% digitonin and then separated on a 3-12% gradient gel as described [9].

Immunoblot analysis of the gel was performed using anti-PKCδ antibody (1:1000) (Abcam, Toronto, ON, Canada) as described [17]. PKCδ was visualized using the Amersham Enhanced Chemiluminescence Western blotting detection system (VWR, Mississauga, ON, Canada). Band intensity was quantified using Image J software. Citrate synthase activity was measured using the citrate synthase assay kit (Sigma-Aldrich, Oakville, ON, Canada, Catalog number CS0720). Data are expressed as mean ± standard deviation of the mean. Comparisons between control and BTHS patient lymphoblasts were performed by unpaired, two-tailed Student *t* test. A probability value of p<0.05 was considered significant.

## Results

The number of each major fatty acyl substituent on individual CL and MLCL molecular species is indicated in **Table 1** and a representative chromatograph of the different CL molecular species between age-matched control and BTHS lymphoblasts is indicated in **Figure 1**. All major molecular species of CL were significantly reduced in BTHS lymphoblasts compared to age-matched control lymphoblasts (**Figure 2A**). Accompanying this reduction in CL molecular species was a general, but not significant increase in most major MLCL species. In contrast, a >20-fold increase (p<0.01) in trioleoyl-MLCL (mass/charge (m/z) 1192) molecular species was observed in BTHS lymphoblasts compared to age-matched control lymphoblasts (**Figure 2B**).

**Table 1.**
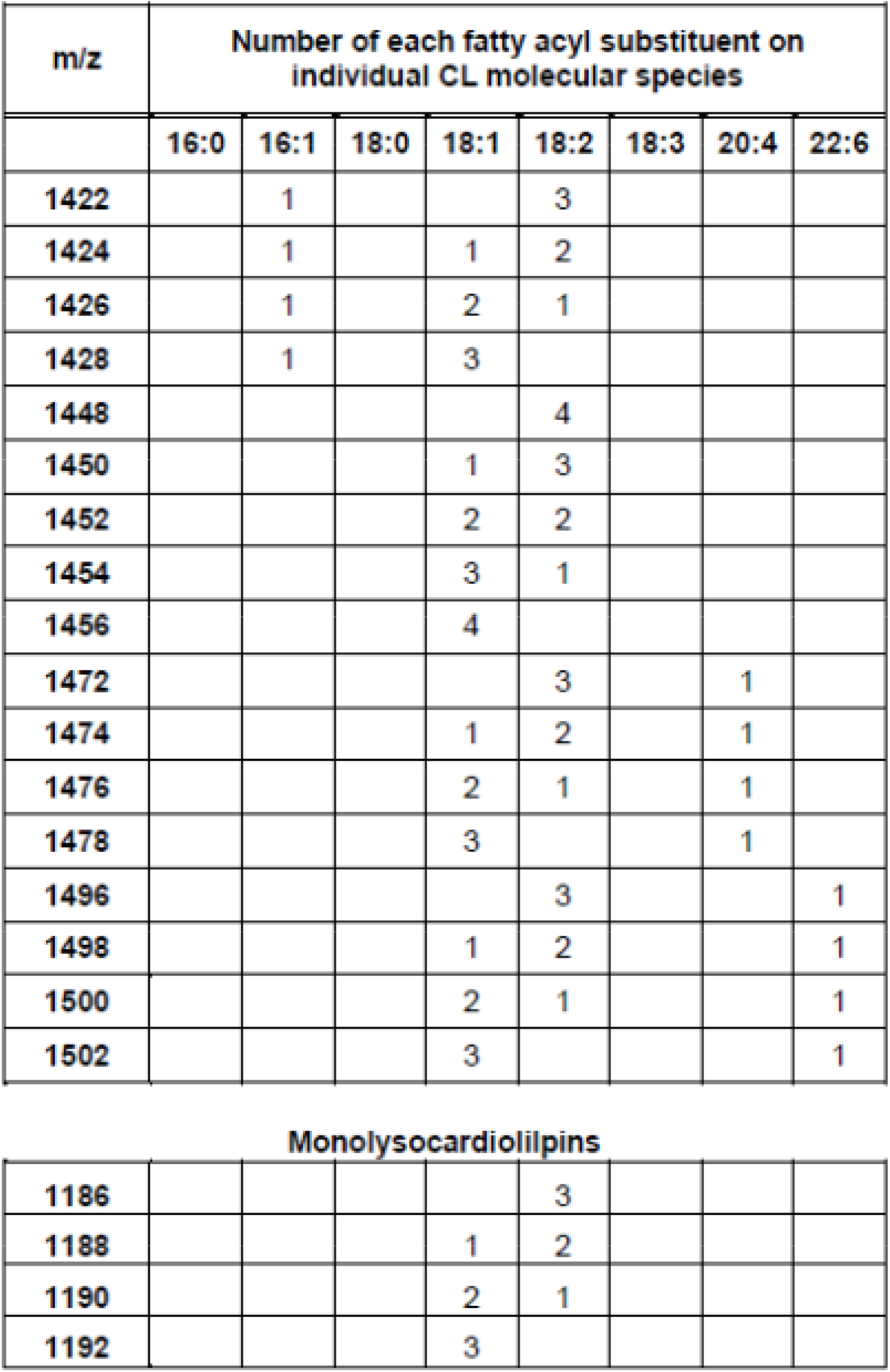
Major CL and MLCL species identified in age-matched control and BTHS lymphoblasts.

| m/z | Number of each fatty acyl substituent on individual CL molecular species |  |  |  |  |  |  |  |
| --- | --- | --- | --- | --- | --- | --- | --- | --- |
|  | 16:0 | 16:1 | 18:0 | 18:1 | 18:2 | 18:3 | 20:4 | 22:6 |
| 1422 |  | 1 |  |  | 3 |  |  |  |
| 1424 |  | 1 |  | 1 | 2 |  |  |  |
| 1426 |  | 1 |  | 2 | 1 |  |  |  |
| 1428 |  | 1 |  | 3 |  |  |  |  |
| 1448 |  |  |  |  | 4 |  |  |  |
| 1450 |  |  |  | 1 | 3 |  |  |  |
| 1452 |  |  |  | 2 | 2 |  |  |  |
| 1454 |  |  |  | 3 | 1 |  |  |  |
| 1456 |  |  |  | 4 |  |  |  |  |
| 1472 |  |  |  |  | 3 |  | 1 |  |
| 1474 |  |  |  | 1 | 2 |  | 1 |  |
| 1476 |  |  |  | 2 | 1 |  | 1 |  |
| 1478 |  |  |  | 3 |  |  | 1 |  |
| 1496 |  |  |  |  | 3 |  |  | 1 |
| 1498 |  |  |  | 1 | 2 |  |  | 1 |
| 1500 |  |  |  | 2 | 1 |  |  | 1 |
| 1502 |  |  |  | 3 |  |  |  | 1 |

**Monolysocardiolipins**
|  |  |  |  |  |  |
| --- | --- | --- | --- | --- | --- |
| 1186 |  |  |  |  | 3 |
| 1188 |  |  |  | 1 | 2 |
| 1190 |  |  |  | 2 | 1 |
| 1192 |  |  |  | 3 |  |

**Figure 1.**
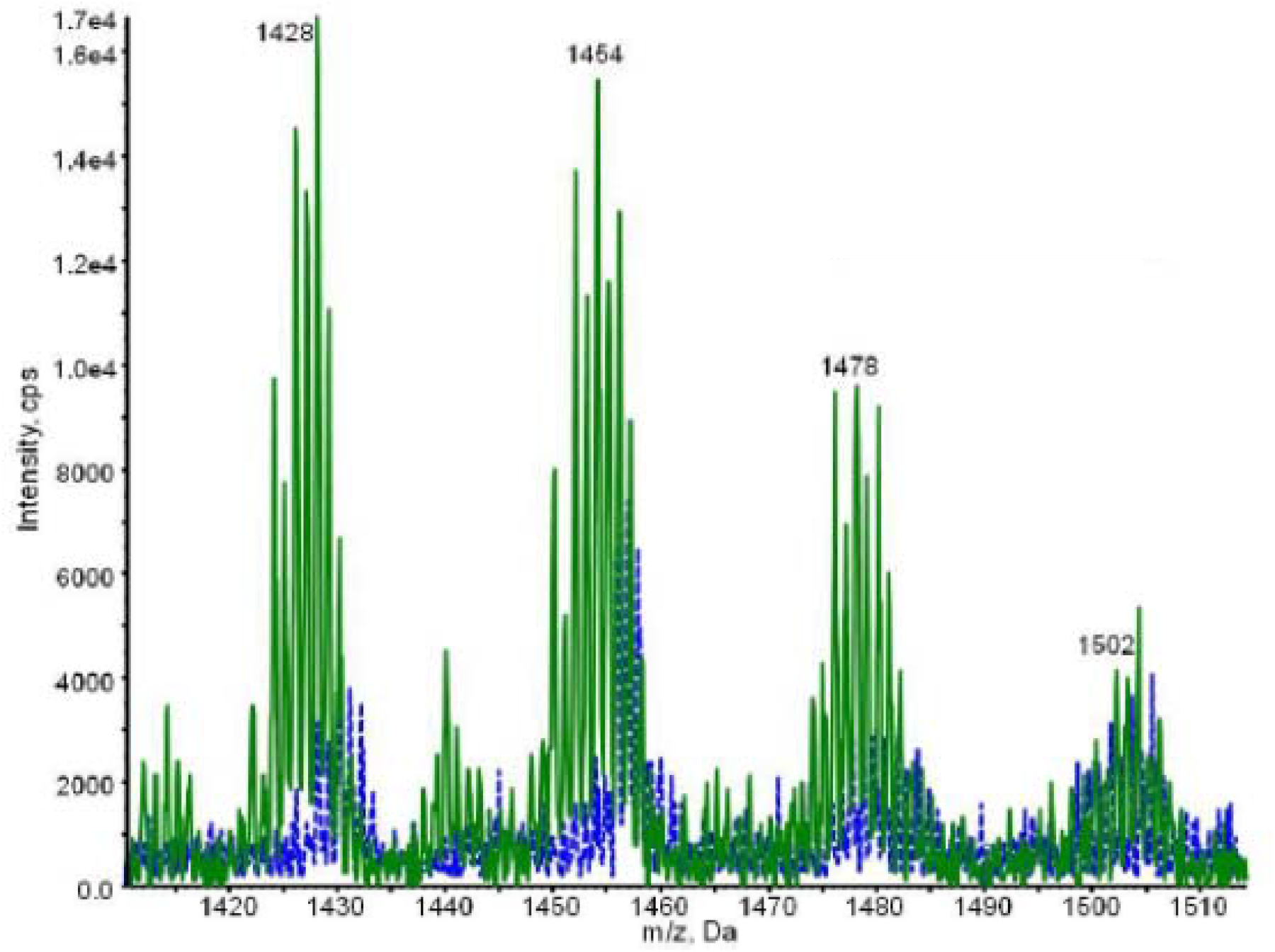
Comparison of CL spectra between age-matched control and BTHS lymphoblasts. Representative chromatograph of the major CL species quantified using ESI-MS coupled with HPLC mass spectrometry of age-matched control (Green solid line) and BTHS lymphoblasts (Blue dashed line).

**Figure 2.**
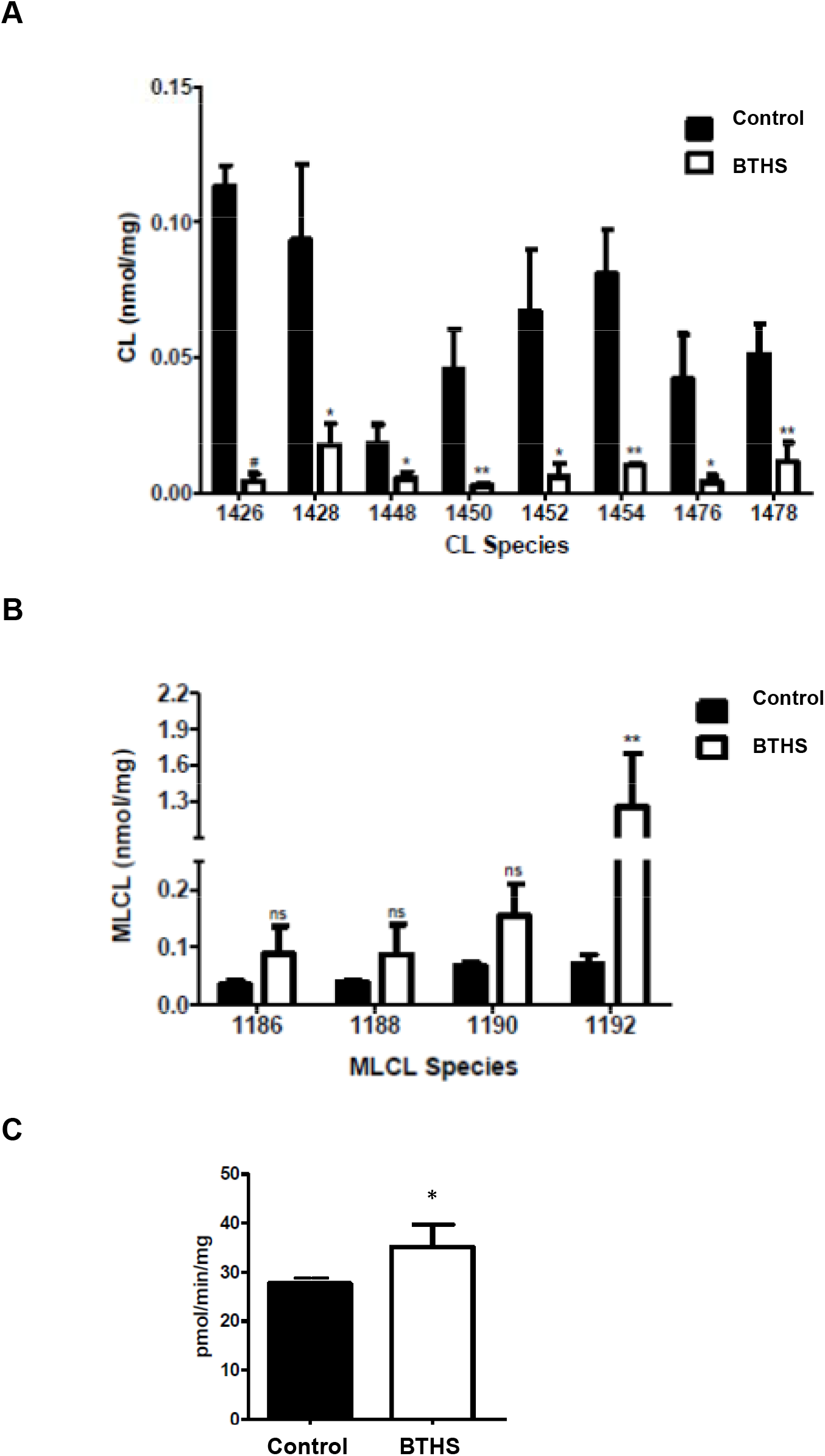
CL levels are reduced and trioleoyl-MLCL and citrate synthase activity elevated in BTHS lymphoblasts. Quantification of the major CL (**A**) and MLCL (**B**) fatty acyl molecular species in age-matched control and BTHS lymphoblasts as described in Materials and Methods. **C**. Mitochondrial fractions were prepared from age-matched control and BTHS lymphoblasts and citrate synthase activity determined as described in Materials and Methods. Data represent the mean ± SD, n=3. #p<0.001, **p<0.01, *p<0.05, ns, not significant.

BTHS patient lymphoblasts exhibited abnormally increased mitochondrial mass [7, 8]. To confirm this, mitochondrial fractions were prepared and citrate synthase activity determined. Citrate synthase activity was elevated 20% (p<0.05) in BTHS lymphoblasts compared to age-matched control cells (**Figure 2C**). Thus, the reduction in CL, increase in MLCL and MLCL/CL ratio and increase in citrate synthase activity was consistent with that observed in BTHS patient B lymphoblast cells.

We previously observed that PKCδ phosphorylation was altered on several sites examined in BTHS lymphoblasts [15]. Since altered phosphorylation may affect PKCδ activation [13], we examined if this was associated with altered PKCδ associated with higher molecular weight complex in mitochondria of BTHS patient lymphoblasts. Mitochondrial fractions were subjected to BN-PAGE followed by immunoblot analysis for determination of PKCδ levels. The two upper bands indicated on the left of the blot are molecular mass markers at 1236 kDa and 1048 kDa, respectively (**Figure 3**). The level of PKCδ located on the gel at a predicted molecular mass near 77.5 kDa was elevated 1.5-fold (p<0.05) in BTHS lymphoblasts compared to age-matched control cells. In contrast, the level of PKCδ located on the gel at approximately 480 kDa was reduced 72% (p<0.01) in BTHS lymphoblasts compared to age-matched control cells. Thus, BTHS lymphoblasts exhibit elevated expression of PKCδ but reduced PKCδ association with a higher molecular weight complex in mitochondria.

**Figure 3.**
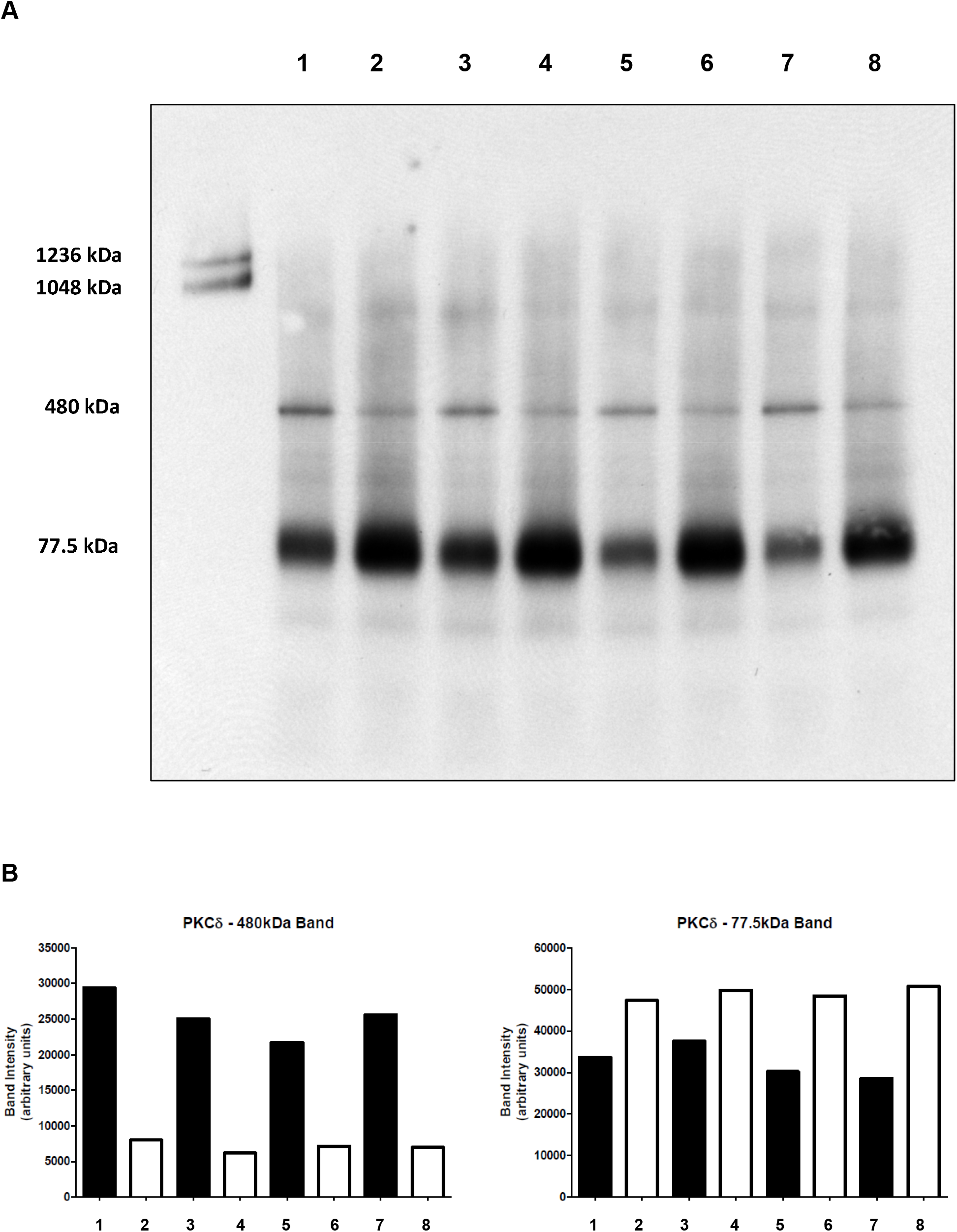
BTHS lymphoblasts exhibit reduced PKCδ association with a higher molecular weight complex in mitochondria. Mitochondrial fractions were prepared from age-matched control and BTHS lymphoblasts and subjected to BN-PAGE followed by immunoblot analysis of PKCδ. **A**. Age matched control (lanes 1, 3, 5 and 7); BTHS lymphoblasts (lanes 2, 4, 6 and 8). Molecular mass markers are in the first lane and indicated on the left. **B**. Quantification of PKCδ.

## Discussion

Protein kinase C delta (PKCδ) is a signaling kinase that regulates a plethora of cellular responses [10-13]. In this study, we examined whether the association of PKCδ with other proteins (which results in the formation of higher molecular weight complexes) was altered in mitochondria of BTHS lymphoblasts. We demonstrate that PKCδ associates with a higher molecular weight complex in lymphoblast mitochondria and that PKCδ higher molecular weight complex formation in mitochondria is reduced in BTHS patient lymphoblasts.

BTHS is a rare X-linked genetic disease and is the only known disease in which the specific biochemical defect is a reduction in CL and accumulation of MLCL [1-3]. We observed a reduction in all major molecular species of CL in BTHS lymphoblasts accompanied by a >20-fold elevation in trioleoyl-MLCL. Previous studies demonstrated an increase in abnormal mitochondrial mass in BTHS patient lymphoblasts [7, 8]. We confirmed this observation in our BTHS patient lymphoblasts through an increase in mitochondrial citrate synthase activity.

BTHS lymphoblasts exhibit elevated oxidative stress and increased reactive oxygen species [8, 9]. Oxidative stress is known to induce expression of PKCδ [18]. We observed increased protein expression of PKCδ in mitochondria of BTHS lymphoblasts compared to controls. The elevated PKCδ levels observed might serve as a compensatory mechanism to increase ATP production in BTHS cells through PKCδ signaling [12]. Additionally, elevated expression of PKCδ promotes mitochondrial proliferation [18]. As indicated above abnormal proliferation of BTHS lymphoblast mitochondria has been previously observed [7, 8]. Phosphorylation of PKCδ is required for its activation [13]. We previously observed alteration in phosphorylation of PKCδ in BTHS lymphoblasts [15]. Altered phosphorylation of PKCδ might contribute to an attenuated mitochondrial PKCδ signaling in these cells.

The molecular components that mediate PKCδ signaling in mitochondria are beginning to emerge. Mitochondria contain a high molecular weight functional complex, which includes cytochrome *c* as the upstream driver of PKCδ, and it uses the adapter protein p66Shc as a platform with vitamin A (retinol) [12, 14, 19]. All four partners are required for functional PKCδ signaling. BN-PAGE immunoblot analysis of mitochondrial proteins not only has the advantage of probing for expression of individual proteins but may additionally be used to detect if these proteins are associated with higher molecular weight complexes. Using this approach we observed reduction in PKCδ association with a higher molecular weight complex in mitochondria.

The PKCδ/retinol complex signals the pyruvate dehydrogenase complex for enhanced flux of pyruvate into the Kreb’s cycle [12, 14]. Interestingly, in the UK BTHS NHS clinic almost half of BTHS boys examined were vitamin A deficient (Nicol Clayton: https://www.youtube.com/watch?v=wNDr_oCTJ7A). However, supplementation with vitamin A did not increase plasma levels. This was not because tissue levels were low but possibly due to increased levels of vitamin A (retinyl esters) in chylomicrons. It is possible that this unique observation is coupled to defective mitochondrial PKCδ signaling which might contribute to a reduced Kreb’s cycle ATP production through alteration in the mitochondrial PKCδ/retinol signaling complex and contribute to the multitude of bioenergetic defects observed in BTHS.

## Acknowledgements

E.M.M. was supported by an MHRC studentship. H.M.Z. is the recipient of a traineeship from the Libyan North American Scholarship Program. G.M.H. is the Canada Research Chair in Molecular Cardiolipin Metabolism. This work was supported by grants from the Heart and Stroke Foundation of Canada, Children’s Hospital Research Institute of Manitoba, the University of Manitoba Research Grants Program and the National Institutes of Health (NIH) USA (P30DK048520).

## Author contributions

E.M.M and G.C.S. performed experiments. H.M.Z. and G.M.H wrote the manuscript. All authors read and edited the manuscript.

## Conflict of interest statement

The authors of this study have no conflict of interest

## Notes

### Competing Interest Statement

The authors have declared no competing interest.

## References

1. Barth PG, Scholte HR, Berden JA, Van der Klei-Van Moorse JMl, Luyt-Houwen IE, Van ‘t Veer-Korthof ET, Van der Harten JJ, Sobotka-Plojhar MA. (1983) An X-linked mitochondrial disease affecting cardiac muscle, skeletal muscle and neutrophil leucocytes. J. Neurol. Sci. 62, 327–355.

2. Kelley RI, Cheatham JP, Clark BJ, Nigro MA, Powell BR, Sherwood GW, Sladky JT, Swisher WP. (1991) X-linked dilated cardiomyopathy with neutropenia, growth retardation, and 3-methylglutaconic aciduria. J. Pediatr. 119, 738–747.

3. Zegallai HM., Hatch GM. (2021) Barth Syndrome: Cardiolipin, cellular pathophysiology, management and novel therapeutic targets. Mol. Cell. Biochem. 476, 605–1629.

4. Xu Y, Malhotra A, Ren M, Schlame M. (2006) The enzymatic function of tafazzin. J. Biol. Chem. 281, 39217–39224.

5. Xu Y, Kelley RI, Blanck TJ, Schlame M. (2003) Remodeling of cardiolipin by phospholipid transacylation. J. Biol. Chem. 278, 51380–5.

6. Valianpour F, Mitsakos V, Schlemmer D, Towbin JA, Taylor JM, Ekert PG, Thorburn DR, Munnich A, Wanders RJA, Barth PG, et al. (2005) Monolysocardiolipins accumulate in Barth syndrome but do not lead to enhanced apoptosis. J. Lipid Res. 46, 1182–95.

7. Xu Y, Sutachan JJ, Plesken H, Kelley RI, Schlame M. (2005) Characterization of lymphoblast mitochondria from patients with Barth syndrome. Lab. Invest. 85, 823–30.

8. Gonzalvez F, D’Aurelio M, Boutant M, Moustapha A, Puech J-P, Landes T, Arnauné-Pelloquin L, Vial G, Taleux N, Slomianny C, et al. (2013) Barth syndrome: cellular compensation of mitochondrial dysfunction and apoptosis inhibition due to changes in cardiolipin remodeling linked to tafazzin (TAZ) gene mutation. Biochim. Biophys. Acta 1832, 1194–206.

9. Mejia EM, Zegallai H, Bouchard ED, Banerji V, Ravandi A, Hatch GM. (2018) Expression of human monolysocardiolipin acyltransferase-1 improves mitochondrial function in Barth syndrome lymphoblasts. J. Biol. Chem. 293, 7564–7577.

10. Reyland ME, Jones DNM. (2016) Multifunctional roles of PKCδ: Opportunities for targeted therapy in human disease. Pharmacol Ther. 165, 1–13.

11. Qvit N, Mochly-Rosen D. (2014) The many hats of protein kinase Cδ: one enzyme with many functions. Biochem. Soc. Trans. 42, 1529–33.

12. Kim YK, Hammerling U. (2020) The mitochondrial PKCδ/retinol signal complex exerts real-time control on energy homeostasis. Biochim. Biophys. Acta Mol. Cell. Biol. Lipids 1865, 158614.

13. Yang Q, Langston JC, Tang Y, Kiani MF, Kilpatrick LE. (2019) The role of tyrosine phosphorylation of protein kinase C delta in infection and inflammation. Int. J. Mol. Sci. 20(6):1498.

14. Acin-Perez R, Hoyos B, Zhao F, Vinogradov V, Fischman DA, Harris RA, Leitges M, Wongsiriroj N, Blaner WS, Manfredi G, et al. (2010) Control of oxidative phosphorylation by vitamin A illuminates a fundamental role in mitochondrial energy homoeostasis. FASEB J. 24, 627–636.

15. Agarwal P, Cole LK, Chandrakumar A, Hauff KD, Ravandi A, Dolinsky VW, Hatch GM. (2018) Phosphokinome analysis of Barth Syndrome lymphoblasts identify novel targets in the pathophysiology of the disease. Int. J. Mol. Sci. 19(7):2026.

16. Sparagna GC, Johnson CA, McCune SA., Moore RL, Murphy RC. (2005) Quantitation of cardiolipin molecular species in spontaneously hypertensive heart failure rats using electrospray ionization mass spectrometry. J. Lipid Res. 46, 1196–204.

17. Chang W, Zhang M, Chen L, Hatch GM. (2017) Berberine Inhibits Oxygen Consumption Rate Independent of Alteration in Cardiolipin Levels in H9c2 Cells. Lipids. 52, 961–967.

18. Lee CF, Chen YC, Liu CY, Wei YH. (2006) Involvement of protein kinase C delta in the alteration of mitochondrial mass in human cells under oxidative stress. Free Radic. Biol. Med. 40, 2136–46.

19. Acin-Perez R, Hoyos B, Gong J, Vinogradov V, Fischman DA, Leitges M, Borhan B, Starkov A, Manfredi G, Hammerling U. (2010) Regulation of intermediary metabolism by the PKCdelta signalosome in mitochondria. FASEB J. 24, 5033–5042.

